# T-cell receptor specific protein language model for prediction and interpretation of epitope binding (ProtLM.TCR)

**DOI:** 10.1101/2022.11.28.518167

**Authors:** Ahmed Essaghir, Nanda Kumar Sathiyamoorthy, Paul Smyth, Adrian Postelnicu, Stefan Ghiviriga, Alexandru Ghita, Anjana Singh, Shruti Kapil, Sanjay Phogat, Gurpreet Singh

**Author notes:** Corresponding author, To whom correspondence should be addressed., Rixensart. RX-60. F1-005, Belgium. Author Contributions AE, GS, PS and SP were involved in the design and/or analysis of the study. GS, AE, PS and NKS acquired the data. AE, AG, AP, AS, GS, NKS, PS, SG and SK analyzed and interpreted the data. All authors were involved in drafting the manuscript or critically revising it for important intellectual content. All authors had full access to the data and approved the manuscript before it was submitted by the corresponding author.

## Abstract

The cellular adaptive immune response relies on epitope recognition by T-cell receptors (TCRs). We used a language model for TCRs (ProtLM.TCR) to predict TCR-epitope binding. This model was pre-trained on a large set of TCR sequences (~62.10^6^) before being fine-tuned to predict TCR-epitope bindings across multiple human leukocyte antigen (HLA) of class-I types. We then tested ProtLM.TCR on a balanced set of binders and non-binders for each epitope, avoiding model shortcuts like HLA categories. We compared pan-HLA versus HLA-specific models, and our results show that while computational prediction of novel TCR-epitope binding probability is feasible, more epitopes and diverse training datasets are required to achieve a better generalized performances in *de novo* epitope binding prediction tasks. We also show that ProtLM.TCR embeddings outperform BLOSUM scores and hand-crafted embeddings. Finally, we have used the LIME framework to examine the interpretability of these predictions.

## Introduction

T Cell Receptors on the surface of a T-cell recognize immunogenic peptides, or epitopes, presented by an HLA molecule on the surface of antigen presenting cells and infected cells. The recognition and interaction of epitopes with TCRs is constrained by the HLA types and the V(D)J gene segment recombination coding for the TCR sequences^1,2^. HLA class I molecules typically present 8-10 amino-acid peptides, which then bind to TCRs with a minimum threshold affinity. These peptides are generated through proteasomal cleavage of intra-cellular proteins and are recognized by CD8 T-cells. HLA class II molecules present larger peptides generated from endocytosed proteins, with variable length exceeding 14 amino-acids. HLA class II-presented peptides are recognized by CD4 T-cells2. TCR-epitope recognition is complex, not only because of the spectrum of physicochemical interactions between the TCR and the HLA-peptide complex, but also due to cross-reactivity: each TCR can recognize many epitopes and each epitope can be recognized by many TCRs^2,3^. This cross-reactivity enables the emergence of public TCRs, i.e., TCRs shared by multiple individuals, and binding to immunodominant epitopes, i.e., recognized by distinct TCRs from distinct individuals^4,5^.

All developments to date indicate that the prediction of TCR-epitope binding predictions is possible, and that reasonable performance can be obtained when predicting binding for unseen CDR3β TCR sequences to epitopes present in the training set. However, a major open question is whether such models perform well on previously unseen epitopes. This has been tested by the authors of ImRex^6^ and Titan models^7^.

Recent developments in natural language processing (NLP) have led to a new paradigm for modeling sequences using pretrained masked language models. By treating amino acids as characters in a language, a self-supervised language model is first trained on the task of predicting the masked characters in a large corpus of protein sequences. This pretrained model can then be finetuned on downstream prediction tasks with limited amounts of labelled data, such as predicting secondary structure or stability^8–10^. This approach is especially relevant for learning TCR-epitope binding where little labelled data is available, and TCR-specific language models (e.g. TCR-BERT) are emerging as a promising direction to solve this task^11^.

The challenges in modelling TCR-epitope binding and the recent achievements in the NLP field applied to proteins motivated us to develop a protein language model for TCR sequences (ProtLM.TCR) and apply it to predict the binding probability between a given TCR and epitope pair. In this work, we focused on TCR CDR3β sequences, HLA class I epitopes and examined the performances of HLA-specific models. We also evaluated the effect of TCR and epitope representability on model performance within the training set, as well as generalization to previously unseen TCRs and epitopes. Further, we compared the performance of TCR language model-based embeddings with other alternatives, such as BLOSUM62 scores, and benchmarked against two publicly available models (Titan and ImRex). Finally, we examined the interpretability of the model predictions using the LIME framework and compared the predicted interactions with resolved 3D structure of peptide HLA-TCR complex.

## Results

### ProtLM.TCR model

We pre-trained a self-supervised masked language model (ProtLM.TCR) on a collection of TCR CDR3β sequences (~62.10^6) from Widrich et al.^12^ and Emerson et al.^13^ by tokenizing the sequences at the character level using a tokenizer with vocabulary of IUPAC amino acid codes.

For self-supervised model training, a random subset of tokens from each sequence was chosen as target labels for the model to predict. Across layers, all other tokens were implicitly considered contextual input. The RoBERTa-style Transformer model processed token indices using self-attention and feedforward modules after converting them to a sum of token and positional vector embeddings^12,14^. The model generates a probability distribution over the token vocabulary for each target token, with the final contextualized token embeddings serving as the prediction head (Figure 1). To train the model, the cross-entropy loss between predictions and target labels was optimized.

**Figure 1:**
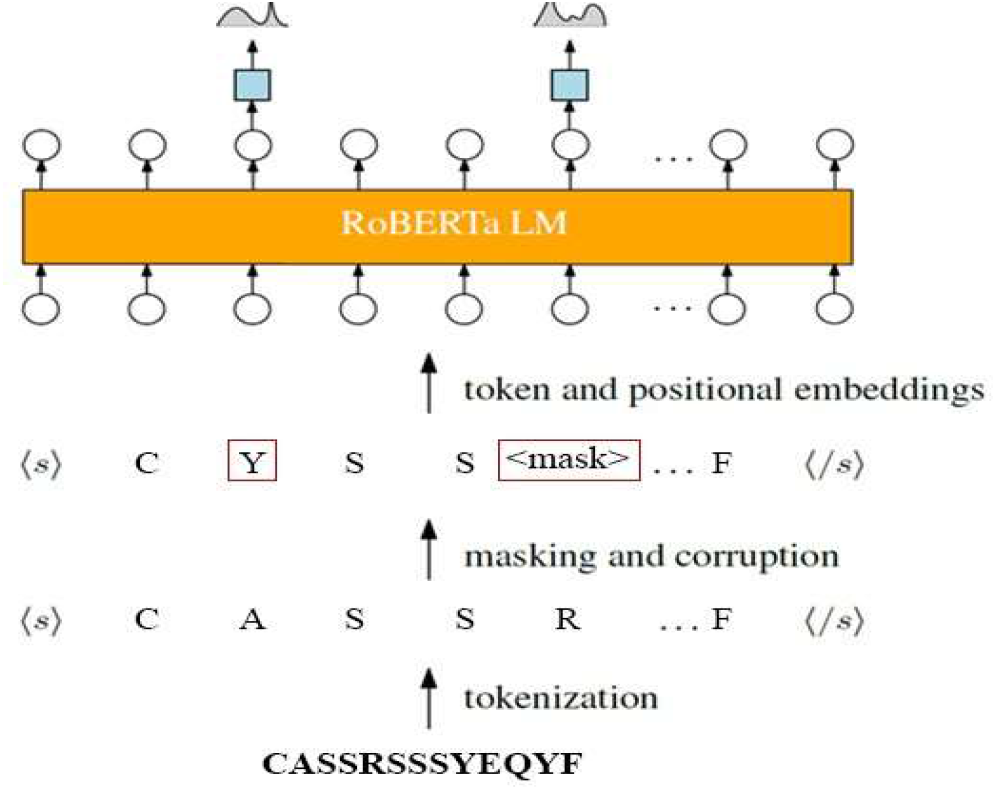
Overview of a masked language modeling pipeline for pretraining RoBERTa-style Transformer language models on large corpora of TCR amino acid sequences. TCR sequences are tokenized at the character level using a predefined amino acid token vocabulary. For each sequence, a random subset of tokens is chosen to serve as training target labels. All other tokens are used for training. Some of this random subsets of tokens are replaced with a mask token, others with a random token, and the remainder are retained. Token indices are converted to a sum of token and positional vector embeddings, which are then processed by alternating layers of self-attention and feedforward modules in the Transformer model. The model generates a probability distribution over the token vocabulary for each token with a target label by acting as a prediction head on the final contextualized token embeddings. By optimizing the cross-entropy loss between the model’s predictions and the target labels, the model is trained.

ProtLM.TCR was fine-tuned for the downstream task of predicting binding between TCR and HLA class I epitope sequences. For this downstream model, we introduced self- and cross-attention mechanisms that take pairs of epitopes and TCR sequences and embed them individually using the pretrained ProtLM.TCR (Figure 2). This downstream cross-attention module consisted of six layers. In each layer the following main processing steps are performed: self-attention (2 layers, one for each entity), cross-attention (2 layers) and feed forward (2 layers). Before each main processing step the inputs are normalized. After normalizing the epitope and TCR sequences separately, they are passed through each of the corresponding six layers, and their results are summed with the original input (i.e., skip connection / residual connection). Finally, after the two feedforward layers the embeddings associated with the <CLS> token are extracted and sent to the fully connected neural network to predict the probability of binding between TCR-Epitope sequences. Importantly, each categorical feature, such as HLA allele, is also converted independently into learnable embeddings. Together, the concatenated representations of sequence and categorical feature embeddings are appended with a dense layer and trained end-to-end to generate binding probabilities for a given TCR and Epitope sequences.

**Figure 2:**
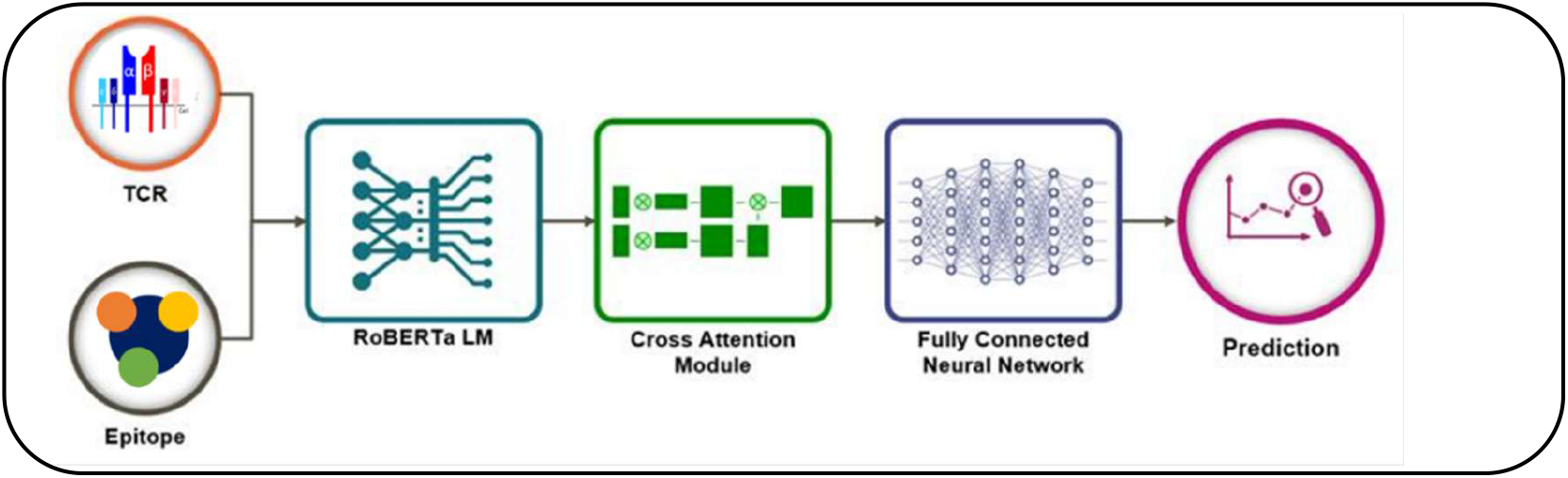
Overview ProtLM.TCR architecture. Using the pretrained ProtLM.TCR, the TCR and epitope sequences are embedded. The downstream cross-attention module has six layers, each processed as follows: to compute self-attention, the epitope and TCR sequences are normalized separately; then cross-attention between the epitope and TCR is computed; finally, two feedforward layers pass TCR and epitope sequences. Moreover, each categorical feature is converted into learnable embedding. A skip connection adds the input to the output of each step to improve training. The resulting multilayer perceptron is trained end-to-end to generate binding probabilities for a given TCR and Epitope sequence.

ProtLM.TCR was applied to predict the probability of binding between TCR-Epitope sequences on the three distinct dataset designs: random assignment of TCR (rTCR), TCR assignment based on protein similarity clusters as computed by ting algorithm (tTCR), Epitope similarity clustering-based assignment (cEPI). Within each of these designs we followed a 5-fold cross validation framework. Here we describe the results for each task and assess the benefit of the additional categorical features such as HLA alleles. The distribution of TCRs, epitopes and their paired instances across these splits is summarized in Table 1.

**Table1:**
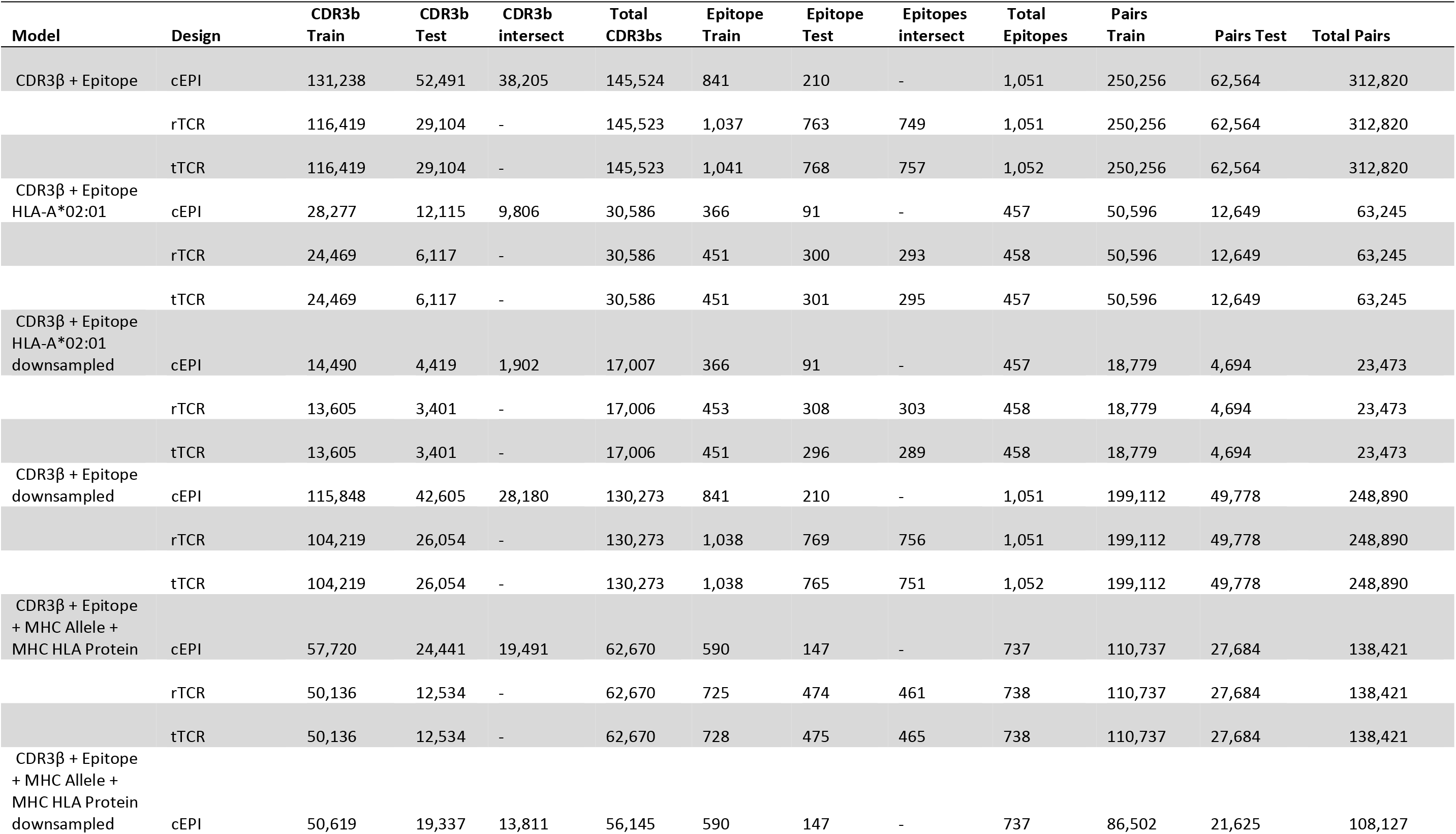
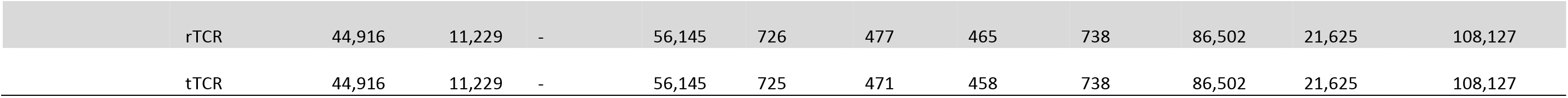
Distribution of TCRs, epitopes and pairs across train and test splits per design.

| Model | Design | CDR3b Train | CDR3b Test | CDR3b intersect | Total CDR3bs | Epitope Train | Epitope Test | Epitopes intersect | Total Epitopes | Pairs Train | Pairs Test | Total Pairs |
| --- | --- | --- | --- | --- | --- | --- | --- | --- | --- | --- | --- | --- |
| CDR3β + Epitope | cEPI | 131,238 | 52,491 | 38,205 | 145,524 | 841 | 210 | - | 1,051 | 250,256 | 62,564 | 312,820 |
|  | rTCR | 116,419 | 29,104 | - | 145,523 | 1,037 | 763 | 749 | 1,051 | 250,256 | 62,564 | 312,820 |
|  | tTCR | 116,419 | 29,104 | - | 145,523 | 1,041 | 768 | 757 | 1,052 | 250,256 | 62,564 | 312,820 |
| CDR3β + Epitope<br>HLA-A*02:01 | cEPI | 28,277 | 12,115 | 9,806 | 30,586 | 366 | 91 | - | 457 | 50,596 | 12,649 | 63,245 |
|  | rTCR | 24,469 | 6,117 | - | 30,586 | 451 | 300 | 293 | 458 | 50,596 | 12,649 | 63,245 |
|  | tTCR | 24,469 | 6,117 | - | 30,586 | 451 | 301 | 295 | 457 | 50,596 | 12,649 | 63,245 |
| CDR3β + Epitope<br>HLA-A*02:01<br>downsampled | cEPI | 14,490 | 4,419 | 1,902 | 17,007 | 366 | 91 | - | 457 | 18,779 | 4,694 | 23,473 |
|  | rTCR | 13,605 | 3,401 | - | 17,006 | 453 | 308 | 303 | 458 | 18,779 | 4,694 | 23,473 |
|  | tTCR | 13,605 | 3,401 | - | 17,006 | 451 | 296 | 289 | 458 | 18,779 | 4,694 | 23,473 |
| CDR3β + Epitope<br>downsampled | cEPI | 115,848 | 42,605 | 28,180 | 130,273 | 841 | 210 | - | 1,051 | 199,112 | 49,778 | 248,890 |
|  | rTCR | 104,219 | 26,054 | - | 130,273 | 1,038 | 769 | 756 | 1,051 | 199,112 | 49,778 | 248,890 |
|  | tTCR | 104,219 | 26,054 | - | 130,273 | 1,038 | 765 | 751 | 1,052 | 199,112 | 49,778 | 248,890 |
| CDR3β + Epitope<br>+ MHC Allele +<br>MHC HLA Protein | cEPI | 57,720 | 24,441 | 19,491 | 62,670 | 590 | 147 | - | 737 | 110,737 | 27,684 | 138,421 |
|  | rTCR | 50,136 | 12,534 | - | 62,670 | 725 | 474 | 461 | 738 | 110,737 | 27,684 | 138,421 |
|  | tTCR | 50,136 | 12,534 | - | 62,670 | 728 | 475 | 465 | 738 | 110,737 | 27,684 | 138,421 |
| CDR3β + Epitope<br>+ MHC Allele +<br>MHC HLA Protein<br>downsampled | cEPI | 50,619 | 19,337 | 13,811 | 56,145 | 590 | 147 | - | 737 | 86,502 | 21,625 | 108,127 |

|  |  |  |  |  |  |  |  |  |  |  |  |
| --- | --- | --- | --- | --- | --- | --- | --- | --- | --- | --- | --- |
| rTCR | 44,916 | 11,229 | - | 56,145 | 726 | 477 | 465 | 738 | 86,502 | 21,625 | 108,127 |
| tTCR | 44,916 | 11,229 | - | 56,145 | 725 | 471 | 458 | 738 | 86,502 | 21,625 | 108,127 |

### ProtLM.TCR model requires only TCR and epitope sequences to capture the TCR-epitope binding prediction signal

We fine-tuned the ProtLM.TCR model end-to-end using TCR CDR3β and their paired epitope amino acid sequences as input (TCR CDR3β + epitope pairs, total: 312,820, Table 1). We followed a 5-fold cross validation random split (total TCRs: 145,524; total epitopes:1,051; Table 1). Our model achieved an average ROC-AUC value of 0.79 for prediction in this task (Figure 3, rTCR). However, we were mindful that random assignment of CDR3β sequences might result in information leakage due to the presence of closely related sequences in the train and test sets (see, rTCR in material and methods). This is especially true for TCRs, which may differ in sequence but share important epitope binding motifs. To eliminate this bias, we pursued an approach that clusters TCRs based on the sequence similarity using the ting algorithm (see, tTCR in material and methods)^15^. Following that, the TCR clusters were divided into 5-fold cross validation train and test sets. In this design, the average ROC-AUC value decreased to 0.71 (Figure 3, tTCR), implying that there was indeed some information shared between train and test sets which might have influenced the performance obtained on random splits. Overall, these findings demonstrate that training ProtLM.TCR models exclusively on CDR3β and epitope sequences was indeed sufficient to capture the signal for TCR-epitope binding prediction to a large extent.

**Figure 3:**
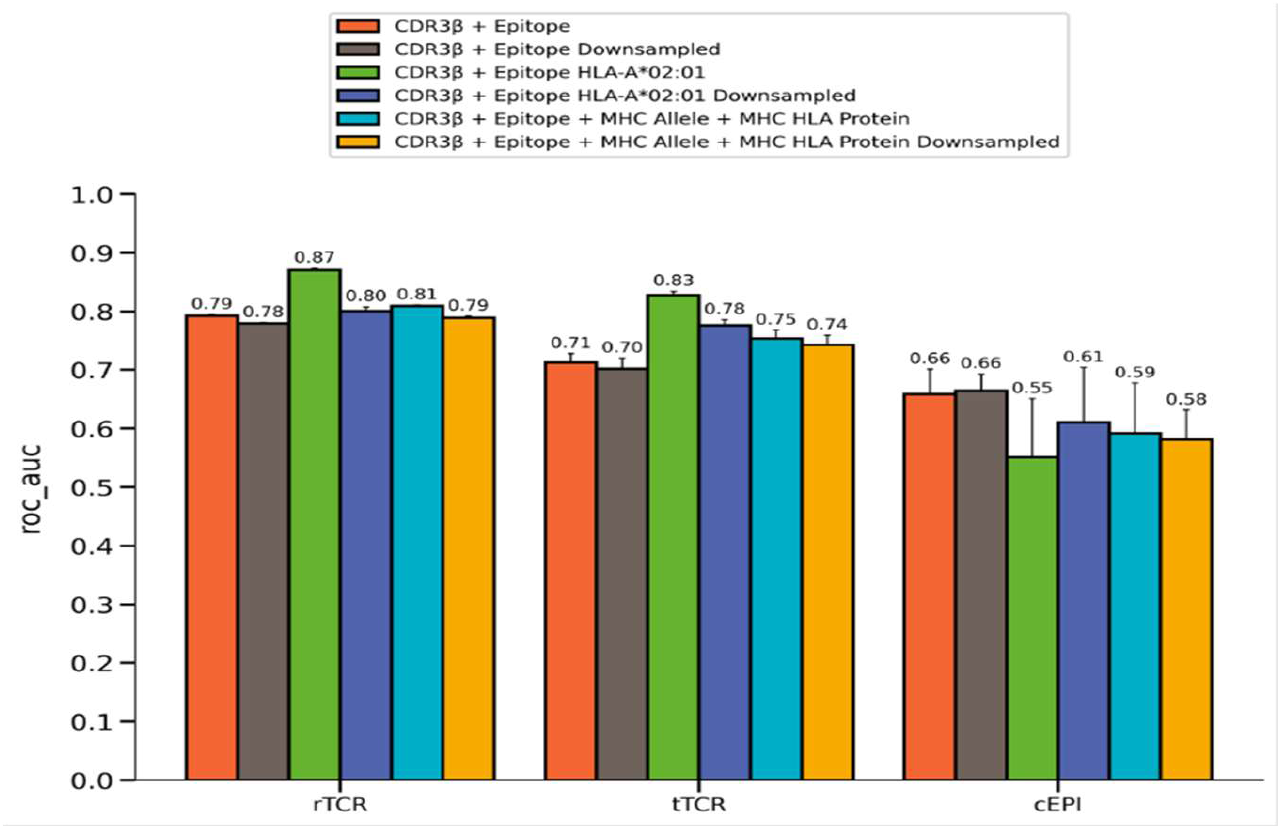
The model performances on three different splitting strategies. We compared the model performances under three different splitting strategies, namely (1) random assignment of TCR (rTCR), (2) TCR assignment based on ting clusters (tTCR), and (3) Epitope clustering-based assignment (cEPI) while ensuring that no epitope clusters are shared between train and test sets. This is the increasing order of complexity for the generalizability of the model. The models were trained and tested using 5-fold cross-validation.

### The predictions made by ProtLM.TCR were robust to the inherent imbalance of the TCR-epitope training datasets

After down-sampling TCRs from the most abundant epitopes, the ROC-AUC values decreased by 1% in both the rTCR and tTCR designs (Figure 3) i.e., 0.79 vs 0.78 (P-value <0.01), and 0.71 vs 0.70 (P-value ~0.26), respectively. This change in performance could be due to a decrease in model overfitting on the most abundant epitope or simply due to a decrease in the number of classes for which the model performed optimally, resulting in a decrease in overall averaged performance. Nonetheless, we conclude, in line with Moris *et al*., that the over-representation of certain epitopes had no discernible effect on model performance^6^.

### Adding HLA information as categorical variable improved the predictions

We noticed that the addition of HLA information as a categorical variable (HLA group and protein) to the basic model (CDR3β + epitope) helped to achieve a marginal increase of 2% for rTCR (0.81 vs 0.79, p-value <0.01; Figure 3) and an increase of ~4% for tTCR (0.75 vs 0.71, p-value <0.01; Figure 3). This improvement in performance demonstrates that including HLA information in the model is beneficial, as expected. Additionally, similar improvements were observed when models were trained with down sampled epitopes to reduce data imbalance; 0.79 vs 0.78 (p-value <0.01) and 0.74 vs 0.70 (p-value <0.01), respectively, for rTCR and tTCR assignments.

### Training the ProtLM.TCR model for specific HLA improved model performances

We sought to further investigate the effect of restricting the task of predicting binding between TCR and epitope for only one type of HLA. We chose to restrict TCR-epitopes to those associated with HLA-A*02:01, the most prevalent HLA type in human populations.

This model achieved the highest ROC-AUCs: 0.87 and 0.83 in rTCR and tTCR assignments, respectively (Figure 3). Down-sampling the epitopes decreased the performances of this HLA-A*02:01 specific model, with a reduction of 7% (0.80 vs 0.87, p-value <0.01) in the rTCR and 5% (0.78 vs 0.83, p-value <0.01) in the tTCR assignment. These findings indicate that while HLA-specific models can outperform pan-HLA model above, however they are more susceptible to data imbalance.

### Generalization to previously unseen epitopes is possible but more difficult due to the training dataset’s limited diversity in the epitope space

We further sought to determine whether our models can generalize to previously unseen epitopes, a more difficult but highly desirable feature. We first clustered epitope sequences (cEPI), then assigned these clusters to train and test sets using 5-fold cross-validation, ensuring that epitope clusters in the test set were not present in the train set. Here, the model trained only on CDR3β, and epitope sequences achieved an average ROC-AUC of 0.66 before and after down-sampling (Figure 3). The model performance did not significantly improve upon addition of HLA covariates (ROC-AUC: 0.59 vs 0.66, p-value ~0.1) and the down-sampling did not improve the results (ROC-AUC: 0.59 vs 0.58, p-value ~0.41). Further, restricting the model to a specific HLA (HLA-A*02:01) negatively affected the generalization (ROC-AUC: 0.55 vs 0.66, p-value <0.05). HLA-specific model generalization did not significantly improve when down-sampling (ROC-AUC: 0.61 vs 0.55, p-value ~0.2). It is worth to know that when adding or restricting features to the model, the numbers of epitopes is reduced significantly (Table 1), which makes these comparisons to the starting model (CDR3β + epitope) approximative.

Overall, these findings indicate that generalization to previously unknown epitopes is possible (66% average ROC-AUC), but better performance may be hampered by the low diversity of the epitope space in current training datasets (maximum:1,052 epitope sequences, Table1).

### TCR language embeddings are better than BLOSUM62 scores

Keeping the architecture of our model the same, we compared different ways to embed TCR and epitope sequences within ProtLM.TCR models (Figure 4). We found that using BLOSUM62 scores as embeddings is significantly decreasing the performances as compared ProtLM embeddings (rTCR and tTCR, Figure 4). Using a specific TCR language for TCR sequences and general protein language for epitope sequences is significantly ameliorating the results for generalization to unseen TCRs (tTCR). ProtLM TCR language embeddings and TCRBert embeddings drive similar performances across all splits, with an amelioration of the results in the rTCR split in favour of TCRBert embeddings.

**Figure 4:**
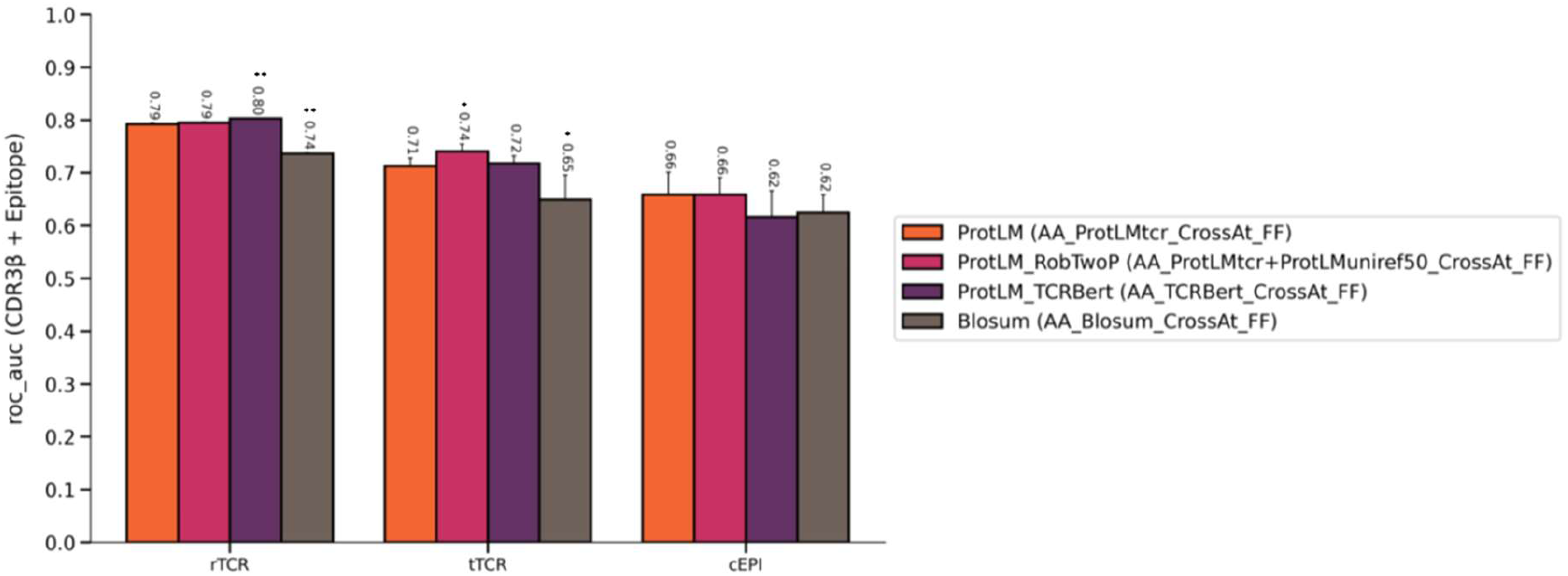
Testing for the impact of embedding strategies on the performances of ProtLM models. We tested four different approaches to embed TCRs and epitopes in ProtLM.TCR models: (i) embedding with ProtLM pretrained TCR language for both TCRs and epitopes (our principal model ProtLM); (ii) embedding TCRs and epitopes using ProtLM pretrained TCR language and ProtLM pretrained language on UNIREF50db (ProtLM_RobTwoP); (iii) embedding both TCR and epitopes using TCRBert pretrained language (ProtLM_TCRBert); and (iv) embedding TCR and epitopes using BLOSUM 62 matrix scores (Blosum). Statistical tests were performed by contrasting each model performances to ProtLM, using Wilcoxon’s test (p-values: *<0.05; **<0.01; ***<0.001).

### Benchmarking against other deep learning-based models for TCR-epitope binding prediction

We benchmarked our models against ImRex and Titan models. We first examined their inference performances in our data held-out splits, by using the models as published by their authors without any re-training (Figure 5A). Both models performed significantly poorly in all the splits showing a limited capabilities for generalization to predictions in new unseen data. However, when retrained, both models (Figure 5B) performed better and approached the performances of ProtLM models, especially in the unseen epitope design (cEPI). We also found that re-training Titan with learnable amino-acid embeddings instead of SMILES for epitopes and BLOSUM62 scores for TCRs ameliorated the results (Figure 5B- Titan, AA retrained versus SMI retrained and finetuned), especially in the unseen TCR design (tTCR).

**Figure 5:**
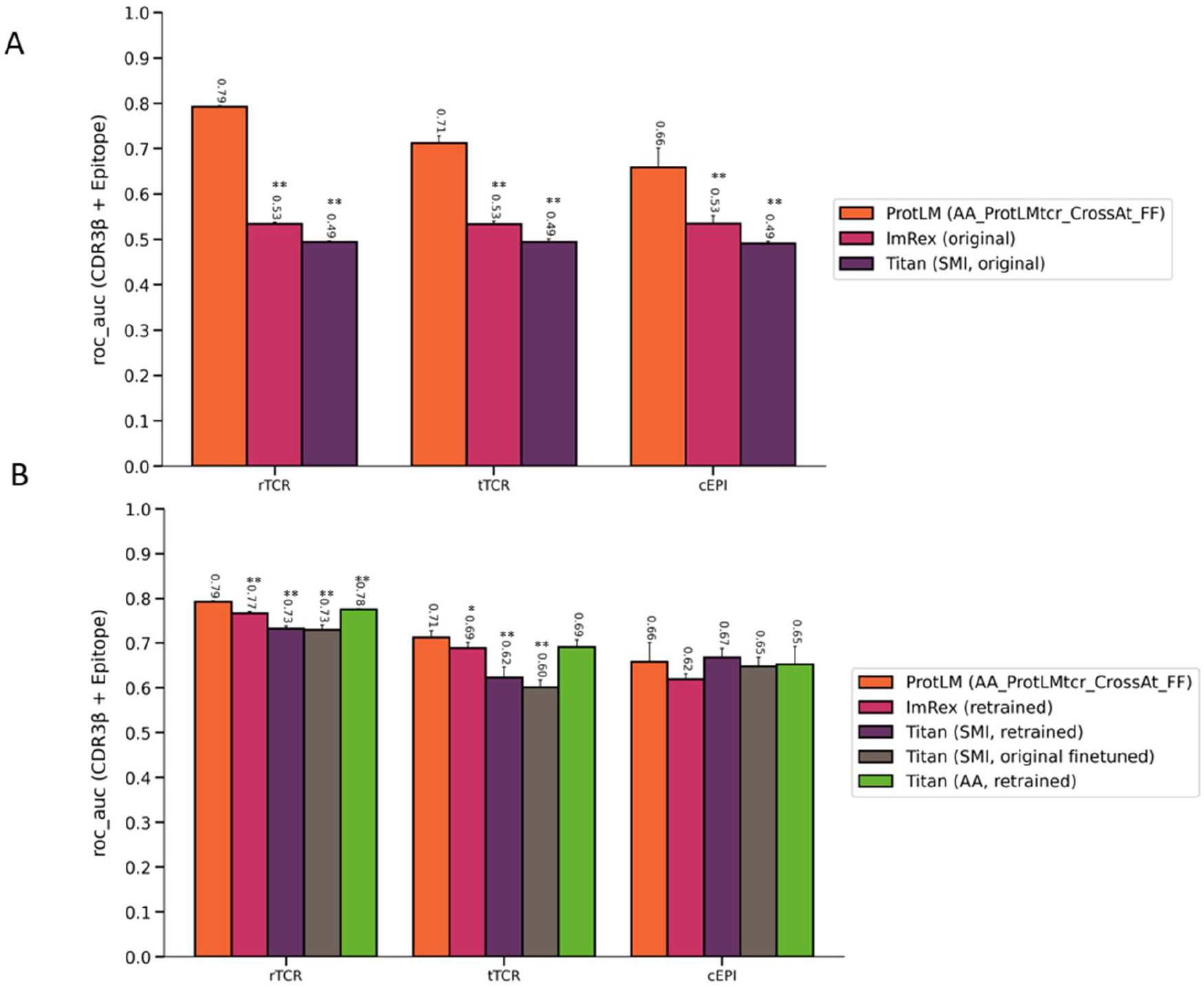
Comparison of the performance of ProtLM.TCR with Titan and ImRex models using our three different splitting strategies. Titan and ImRex model performances were evaluated using the same data splits used to train and evaluate ProtLM.TCR models. **A)** We used the original models exclusively for inference where the models were used exactly as they were obtained from their respective repositories and tested on the held-out data split. **B**) All models were trained from scratch for the retrained comparisons. For Titan, we used SMILES (SMI) or amino-acid (AA) encodings for epitope sequences; TCR sequences are always encoded as AA; we either fine-tuned the models using their original embeddings (BLOSUM) or completely retrained them. Statistical tests were performed by contrasting each model performances to ProtLM, using Wilcoxon’s test (p-values: *<0.05; **<0.01; ***<0.001).

ImRex models embed amino acids using physicochemical feature engineering. ProtLM versus ImRex results (Figure 5B) suggest that the TCR language might have captured beyond the physicochemical properties of the amino acids involved in binding between TCR-epitope sequences.

In conclusion, the results demonstrate that TCR-epitope prediction is learnable, that protein language modeling eliminates the need to explicitly feature-engineer the physicochemical properties of amino acids, and that additional data would need to be generated on the TCR and epitope binding to achieve generalization performances greater than 0.66 ROC-AUC for predicting the binding of novel epitopes.

### LIME framework can be used to explain the TCR-epitope binding predictions by ProtLM.TCR models

We used the Local Interpretable Model-Agnostic Explanations (LIME) framework to explain TCR-epitope binding predictions made by ProtLM.TCR models. We illustrate our approach by interpreting the binding predictions using the TCR and epitope sequences in the resolved pHLA-TCR-epitope complex for the PDB entry: 2VLJ^16^. This complex shows the interaction of CASSRSSSYEQYF CDR3β with its cognate GILGFVFTL epitope as presented by HLA-A*02:01 (Figure 6A). We calculated the pairwise amino acid interactions between CDR3 β and epitope sequences using this PDB structure to highlight the closest distances showing H-bond interactions (Figure 6B). Interestingly, we were able to deduce the importance of Arginine (R) and Serine (S) residues in positions 6 and 7, respectively, in driving the CDR3 β interactions with this epitope from the interpretation of the LIME scores. Additionally, LIME emphasized the significance of Serine at positions 4 and 8, as well as Tyrosine(Y) and Glutamine(Q) at positions 9 and 11, respectively (Figure 6C). By comparing the LIME heatmap (Figure 6C) to the distance matrix table (Figure 6B), we discovered that the highlighted amino-acid positions relevant for the interaction had a high degree of overlap. This was particularly interesting because, even though ProtLM.TCR was trained using only linear sequences, we were still able to identify specific amino acids (R6 and S7) as contributing to the interaction via LIME analysis (Figure 6D), which could also be visualized using the PDB 3D interactions.

**Figure 6:**
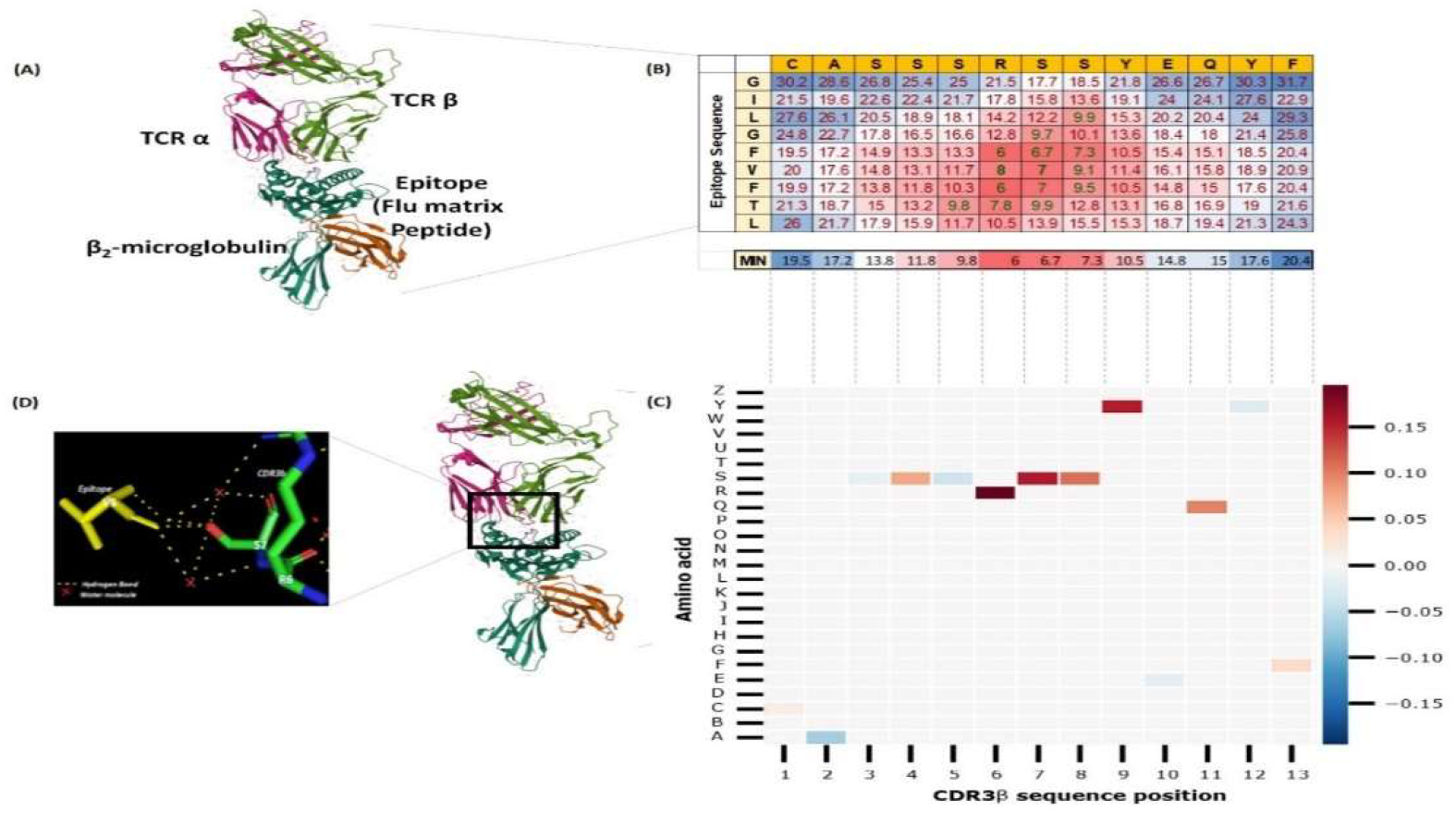
Local Interpretable Model-Agnostic Explanations (LIME) for The binding between TCR and epitope sequence for PDB ID 2VLJ. (A) 3D structure of the 2VLJ entry in PDB database, which shows a TCR beta (CASSRSSSYEQYF): TCR alpha complex binding GILGFVFTL epitope presented by HLA class I (HLA-A*02:01):Beta-2 macroglobulin complex (B) Heatmap visualizing the distance matrix (in Å units) between amino acids in TCR CDR3β to their pairs in the epitope side. Distances less than 8 Å showed strong H-bond interactions (e.g., V6 epitope- R6 and S7 CDR3β interactions showed 8 Å and 7 Å respectively). The smaller the distance, the higher the interaction. The below row (MIN) represents the minimum distance of CDR3β to each of the epitope amino acids. (C) Heatmap visualization showing the LIME scores as computed for each amino acid in the CDR3β (CASSRSSSYEQYF) when predicting its interaction with GILGVFTL epitope. The position of each amino acid in the sequence is presented in the x-axis and its identity in the y-axis. Red color represents strong effect in the binding predictions, whereas blue represents weak effect. (D) The crystal structure of the complex shows the interactions of epitope V6 amino-acid with CDR3β R6 and S7 amino-acids via hydrogen bond and water molecules.

We have checked two more LIME interpretations of TCR-epitope bindings in PDB IDs 3O4L and 3GSN (Supplementary Figures S1 and S2). In the case of 3O4L, we found the importance of Arginine (R), Glycine (G), Tyrosine (Y) and Threonine (T) at position 4, 8, 11 and 7 whereas in 3GSN, Threonine (T) at position 7 respectively. These amino acids are responsible for CDR3 β interactions with their cognate epitope. These highlighted amino acids were further confirmed by visualization of H-Bonds in PDB 3D interactions.

## Discussion

Accurate predictions of T-cell receptor (TCR)-epitope binding will facilitate the prioritization and rationalization of vaccine antigens, as well as other biomedical applications. However, developing such models is complicated by the scarcity of data, particularly on the epitope side, and the complexity of the biological problem. Indeed, the diversity of TCRs, HLAs, and their restricted epitopes, combined with cross-reactivity, complicates the task of TCR-binding prediction. The classical immunoinformatic tools do not take T-cell receptors into account and rely heavily on HLA-peptide binding affinity prediction, which does not guarantee that the predicted epitopes are immunogenic (i.e., will bind to TCRs), and thus the achieved performance is prone to high rates of false positives, particularly when testing for immunogenicity^17,18^. Recently, pan-HLA models have emerged that predict the TCR-epitope bindings directly from their respective amino acid sequences. Among the challenges in developing such models are the following: (i) how to embed amino acids into vectors suitable for training machine/deep learning models while preserving their conservation and physicochemical properties? (ii) how to create unbiased datasets from published TCR-epitope datasets that account for HLA specificity, cross-reactivity, and a bias toward positive binders? and (iii) how to interpret the model predictions and identify key amino acid interactions?

Our approach alleviates these difficulties by developing a protein language model for TCR (ProtLM.TCR) using a large corpus of CDR3β TCR sequences, similarly to previously published protein language models, for which the embeddings have been shown able to capture the physicochemical properties of amino acids and their probabilistic co-occurrences as shaped by evolution^14,19^. The learned embeddings are then used in fine-tuning models to predict the binding between a given epitope and TCR sequences. These models were trained on a TCR-epitope dataset prepared from published instances by balancing negative and positive binders across HLA types. Our results indicate that training models on only CDR3β and epitope sequences was sufficient to capture most of the binding prediction information. When HLA covariates were included, performance was slightly improved. However, when such information was included, the data size decreased, which may have skewed the comparisons. We anticipate that if the model is trained with the same amount of data but with HLA and TCR-α information included, performance will improve significantly.

By restricting the model to a single HLA (HLA*A:02), the results improved, indicating that HLA-specific models may be more accurate than pan-HLA models. The fact that this HLA*A:02 has more data, however, may reflect a data-availability bias. With additional data being generated for additional HLAs, follow-up studies should address this issue.

Our models were initially trained and tested on random splits (rTCR); however, equivalent performance was obtained by controlling TCR sequences across train and test data splits in clusters that shared not only similar sequences but also motifs (tTCR). This generalization to previously unseen TCRs in the last TCR-clustered data splits achieved reasonable results, demonstrating that TCR-epitope predictions are learnable and can be used to screen new TCRs for binding to the training set epitopes.

Further, by clustering epitope sequences (cEPI), we controlled epitope similarity between train and test splits. While generalization to unknown epitopes performed poorly, the ROC-AUC metrics were significantly greater than random expectation. Taken together, these findings demonstrate that the TCR-epitope binding prediction is both learnable and generalizable to novel epitopes, though the scarcity of epitope sequences in the training dataset may act as a constraint on improved generalization.

We showed that TCR language models eliminate the need for BLOSUM scores and feature engineered embeddings of amino acids when the embeddings are not learned as part of the model training. Additionally, our approach of balancing positive and negative binders across HLA types aided Titan and ImRex models in learning more effectively, as evidenced by their improved performances when trained on our data splits compared to the inference-only performances of their published versions. Our findings imply that any model capable of learning motifs (e.g., deep learning and nonlinear models) can be used if the upstream amino-acid embeddings accurately represent their physicochemical properties and occurrences, in line with recent observations by Wu et. Al.^11^. Finally, we examined the model using the Local Interpretable Model-Agnostic Explanations (LIME) technique to gain insight into the putative driving interactions between CDR3β and the epitope sequences is captured. We further inspected these interpretations by comparing the putative TCR-epitope interaction sites, identified as significant amino acids via LIME, to the experimentally determined 3D structures from PDB. We demonstrated that our model had learned amino acid interactions that are likely to be involved in TCR-epitope binding based on their physical proximity as determined by the 3D structure. While caution should be exercised with all such interpretations, as LIME will always provide an interpretation that may or may not agree with the experimental data, this approach provides an intuitive means of quickly verifying the model predictions. In future studies, we will aim to conduct a comprehensive study of these aspects.

## Conclusions

We have developed models for predicting and interpreting binding between T-cell receptors (CDR3β) and HLA class I epitopes using Protein Language Modeling, and we have shown that this provides a better approach for embeddings of TCR and epitope amino acid sequences. We have also generated a standard training and evaluation dataset and compared our model performance to those previously published models. Doing this, we have achieved decent accuracy in predicting the binding of previously unseen TCRs and epitopes. To aid researchers in deciphering the antigen-specific landscape and underlying immune responses in a variety of disease-related studies, we have also illustrated the model’s understanding of the interactions between TCRs and the relevant epitope sequences using LIME. Lastly, we also stress the critical nature of increasing data generation in this field, particularly in epitope space, to improve accuracy and generalization. This is especially critical for developing pan-HLA models.

## Material and Methods

### Data Preprocessing

We gathered the published human TCR-epitope pairs from multiple public databases: VDJdb, IEDB, McPAS-TCR and PIRD^18,20–22^. These datasets were then merged and cleaned to remove redundant samples (see Appendix 1). Negative samples were generated by randomly pairing TCR sequences (CDR3βs) with non-binding epitope-HLA complexes. Thereafter, we clustered the TCR sequences using ting and the epitope sequences using ImmunomeBrowser^16,23^. To account for the epitope imbalance, we downsampled the datasets by imposing a limit on the number of TCRs per epitope. Finally, we created three distinct versions of the dataset:

1. random assignment of TCRs (**rTCR**),
2. TCR assignment based on ting clusters (**tTCR**)
3. Epitope clustering-based assignment (**cEPI**)

In the latter case we ensured that no epitope clusters are shared between train and test sets. For cross-validation purposes, the dataset was divided into five-folds for each of these versions. This distinction in information overlaps between the three versions enabled a more refined assessment of the model’s generalizability. Table 1 contains the total number of TCRs, epitopes, and HLA categorical variables, as well as their distribution by train and test splits.

### Benchmarking

ImRex and TITAN models were trained on the same dataset of TCR (CDR3β) and epitope sequences that were used to train our model. These comparisons used the same 5-fold cross-validation splits and three categories of assignment as described above for our model i.e., rTCR, tTCR, and cEPI. We used the average ROC-AUC and 95 percent confidence interval obtained after evaluating the test splits as an evaluation metric. Additional information is included in Appendix I.

### LIME and interpretability

The LIME algorithm was used to determine the importance of each amino acid in the interacting amino acids^24^. To interpret the significance of the amino acids involved, a given input sequence was subjected to a masking process in which a fixed-size local dataset was created by generating “random” perturbations from the given sequence, and then a ridge regression linear model was trained on the resulting dataset to generate scores that were then normalized with L1-norm to generate the final scores. Amino acids with positive LIME scores support the prediction, whereas amino acids with negative scores contradict the prediction. This means that positively scored amino acids are likely to be the primary drivers of binding.

For the analysis presented in this paper, the Protein Data Bank (PDB) was used to download crystal structures^25^. To determine the interaction between amino acids for a chosen pair of CDR3β and epitope sequences, we calculated the residue distance matrix using the PyMOL tool^26^. The computed distances from maximum to minimum were depicted using a blue to red color scheme. The strongly interacting amino acids between CDR3β and epitope were visually examined using PyMOL computed hydrogen bond interactions.

## Sponsorship and Acknowledgements

This work was sponsored by GlaxoSmithKline Biologicals SA which was involved in all stages of the study conduct and analysis. We acknowledge and thank Matthias Bal for his involvement in the initial stages of this project. We thank Alex Pysik, Amin Khan, Yannick Vanloubbeeck, Fernando Ulloa-Montoya, Guglielmo Roma, Janet Effler, Kalyani Anantapantula, Laurent Sorber, Ludmila Tydlitátová, Marie Toussaint, Michael Ferenczy, Nicolas De Neuter, Rebecca Stephens, Roberto Spreafico, Sai Jasti, Tanguy Naets, Thomas Peel, and Viviane Bechtold for their support and valuable inputs.

## Appendix I

## Data Preprocessing

- Data for this study was extracted from VDJdb, IEDB, McPAS-TCR and PIRD databases and filtered for human HLA class I.
- Basic data cleansing was performed, and HLA categories were standardized as per WHO nomenclature.
- It was ensured that the CDR3β sequences have always the conserved C and F amino acids at N- and C-terminal respectively by explicitly adding the caps wherever necessary.
- Dataset was then filtered based on the minimum length of sequences.
- Samples with non-standard amino acids, undetermined HLA class were removed.
- Samples duplicated between different data sources were identified and removed.
  - TCR sequences were clustered using ting^28^ under default settings with reference dataset: gliph-1.0/db/TCRab-naive-refdb-pseudovdjfasta.fa and kmer file: LULO-TC-DN6_Tc_N_CD40L-.tsv
- epitope sequences were clustered using IEDB clustering tool^25^ with 70% minimum sequence identity threshold and the recommended clustering method: cluster-break for clear representative sequence.
- Negative sample generation:
  ○ For each epitope, all the TCRs from TCR clusters that are not known to bind that epitope were selected as non-binding candidates for that epitope.
  ○ Negative samples are created by replacing the binding TCRs in the positives, with randomly sampled non-binding candidates to achieve a 1:1 target ratio for each epitope, while always keeping the HLA constraint.
- Six different versions of the dataset were created:
  ○ v1: CDR3β and epitope as input features
  ○ v2: CDR3β, epitope and HLA categories as input features
  ○ v3: same as v1, but restricted to HLA-A*02:01
  ○ v4: down sampled version of v1.
  ○ v5: down sampled version of v2.
  ○ v6: down sampled version of v3.
  ○ These six datasets were split using the below strategies:
  ○ Random splits: 5-fold cv such that unique CDR3β sequences are randomly split between folds
  ○ ting TCR clusters-based splits: 5-fold cv such that TCR clusters are split independently between folds i.e., a given cluster can only be bucketed in one of the folds.
  ○ epitope-clusters based splits: 5-fold cv such that epitope clusters are split independently between folds

## Benchmarking

1. ImRex:
  - Model: Architecture 0 - 2020-07-30_11-30-27_trbmhci-shuffle-padded-b32-lre4-reg001 – a model trained on VDjdb (august 2019 release).
  - Hyper parameter changes: batch size 32 -> 128, dropout_conv: 0.25->0.1; max_length_epitope 11->45; max_length_CDR3β 20->40; lr 0.0001 -> 0.00015; regularization 0.1 ->0.008; number of train epochs:20; We used the provided hand-crafted amino-acid physico-chemical feature. These parameters were identified by doing hyperparameter search on one of our data splits (the ting model_0 split_0_train.csv subset).
  - Code version: origin/master from 24 Feb 2021.
2. Titan: We trained 3 architectures using TCR CDR3β and Epitope sequences. The configuration was downloaded from IBM.box.com/v/titan_dataset
  - Architecture 0 – Inference: trained and published model from IBM.box.com/v/titan_dataset, where TCR CDR3β amino-acids are encoded as amino-acids and embedded using BLOSUM, and epitope sequences are encoded using SMILES and the embedding were learned.
  - Architecture 1 – Titan retrained: TCR CDR3β are encoded as amino-acids and embedded using BLOSUM and epitope sequences are encoded using SMILES and the embeddings are learned.
  - Architecture 2 – Titan finetuned: - finetuning the model provided in IBM.box.com/v/titan_dataset that was trained by the authors on full TCR proteins encoded as AA and embedded using BLOSUM, and epitopes encoded as SMILES and embeddings are learned. Here we used TCR CDR3β amino-acids and embeddings using BLOSUM and epitopes using SMILES and embeddings are learned. We changed the number of epochs to 20.
  - Architecture 3 – Titan retrained: TCR CDR3β and epitopes are encoded as AA and embeddings are learned.
  - Code version: origin/main from 16 Sep 2021

---

**Figure S1:**
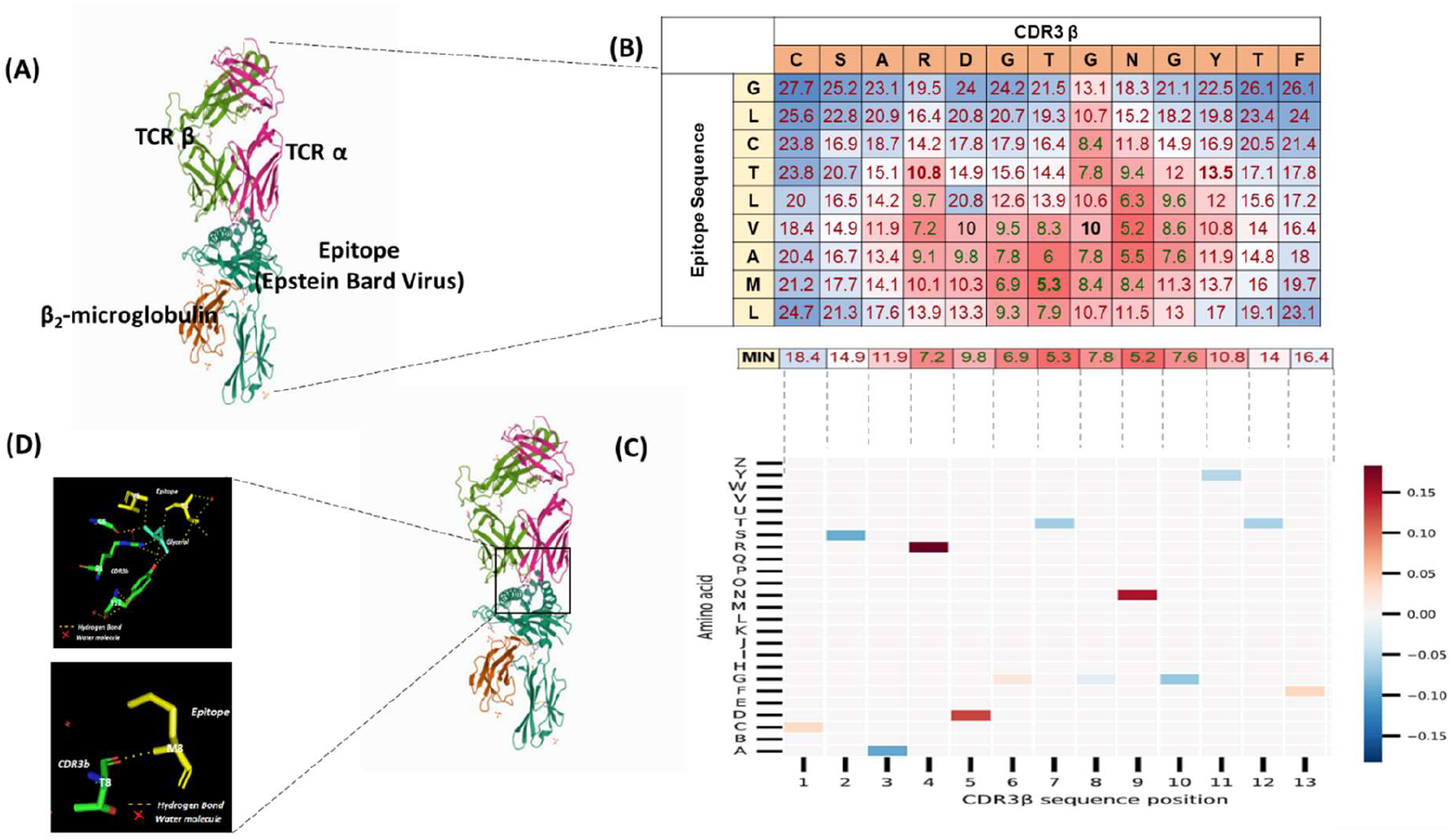
Local Interpretable Model-Agnostic Explanations (LIME) for the binding between TCR and epitope sequence for PDB ID 3O4L. (A) 3D structure of the 3O4L entry in PDB database, which shows a TCR beta (CSARDGTGNGYTF): TCR alpha complex binding GLCTLVAM epitope presented by HLA class I (HLA-A*02:01):Beta-2 microglobulin complex (B) Heatmap visualizing the distance matrix (in Å units) between amino acids in TCR CDR3β to their pairs in the epitope side. Distances less than 8 Å showed strong H-bond interactions. The smaller the distance, the higher the interaction. The below row (MIN) represents the minimum distance of CDR3β amino acids to each of the epitope amino acids. (C) Heatmap visualization showing the LIME scores as computed for each amino acid in the CDR3β chain (CSARDGTGNGYTF) when predicting its interaction with GLCTLVAM epitope. The position of each amino acid in the sequence is presented in the x-axis and its identity in the y-axis. Red color represents strong effect in the binding predictions, whereas blue represents weak effect. (D) The crystal structure of the complex showing the interactions of epitope T4 and V6 amino-acid with CDR3β R4/Y11 and G8 amino-acids form hydrogen bond via Glycerol whereas, T7 of CDR3 β and M8 epitope form direct H-bonds.

**Figure S2:**
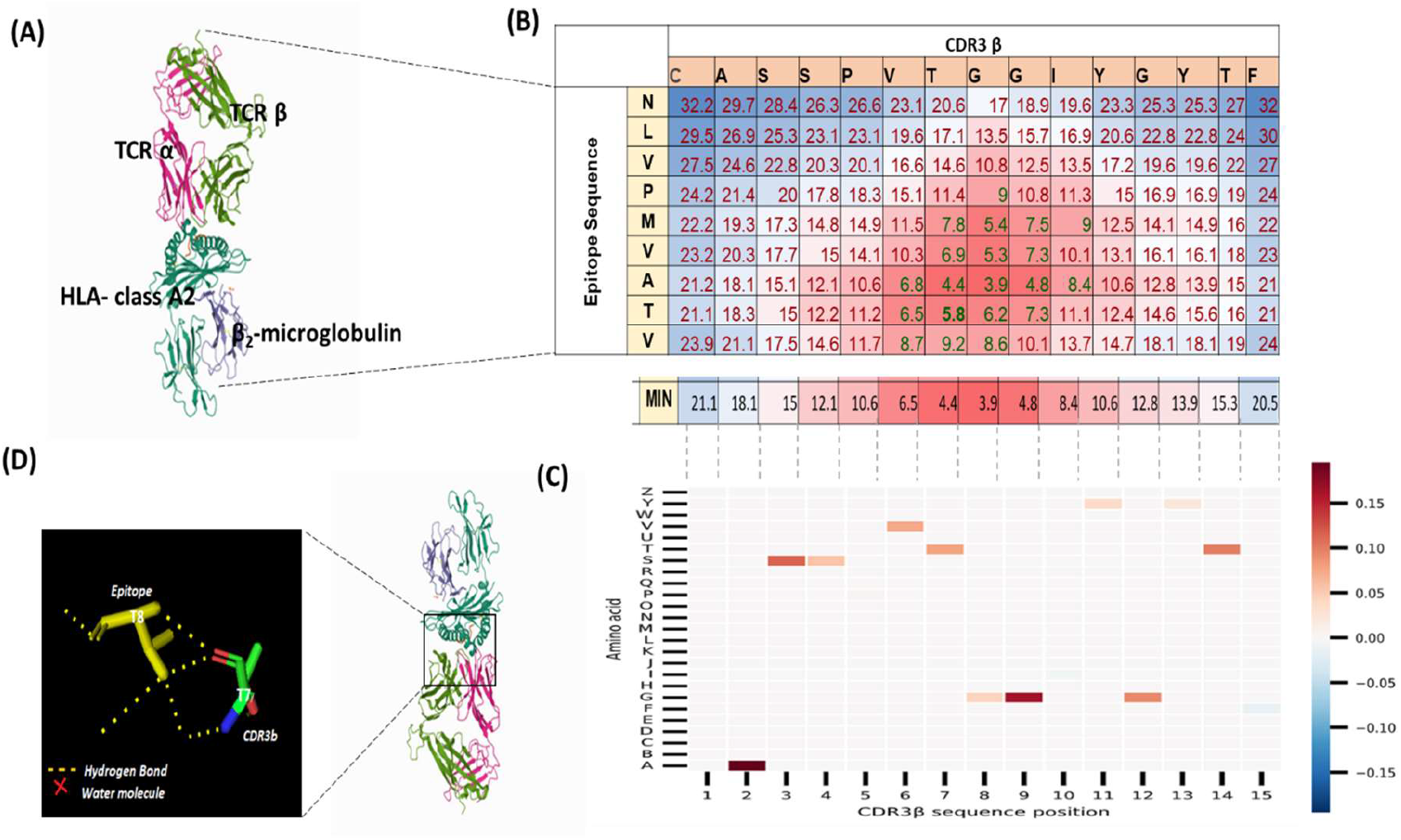
Local Interpretable Model-Agnostic Explanations (LIME) for the binding between TCR and epitope sequence for PDB ID 3GSN. (A) 3D structure of the 3GSN entry in PDB database, which shows a TCR beta (CASSPVTGGIYGYTF): TCR alpha complex binding NLVPMVATV epitope presented by HLA class I (HLA-A*02:01):Beta-2 microglobulin complex (B) Heatmap visualizing the distance matrix (in Å units) between amino acids in TCR CDR3β to their pairs in the epitope side. Distances less than 8 Å showed strong H-bond interactions. The smaller the distance, the higher the interaction. The below row (MIN) represents the minimum distance of CDR3β amino acids to each of the epitope amino acids. (C) Heatmap visualization showing the LIME scores as computed for each amino acid in the CDR3β chain (CASSPVTGGIYGYTF) when predicting its interaction with NLVPMVATV epitope. The position of each amino acid in the sequence is presented in the x-axis and its identity in the y-axis. Red color represents strong effect in the binding predictions, whereas blue represents weak effect. (D) The crystal structure of the complex showing the interactions of epitope T8 with T7 of CDR3β via H-bonds.

